# Teleoperation of an ankle-foot prosthesis with a wrist exoskeleton

**DOI:** 10.1101/2020.07.17.209049

**Authors:** Cara G. Welker, Vincent L. Chiu, Alexandra S. Voloshina, Steven H. Collins, Allison M. Okamura

## Abstract

**Objective:** We aimed to develop a system for people with amputation that non-invasively restores missing control and sensory information for an ankle-foot prosthesis.

**Methods:** In our approach, a wrist exoskeleton allows people with amputation to control and receive feedback from their prosthetic ankle via teleoperation. We implemented two control schemes: position control with haptic feedback of ankle torque at the wrist; and torque control that allows the user to modify a baseline torque profile by moving their wrist against a virtual spring. We measured tracking error and frequency response for the ankle-foot prosthesis and the wrist exoskeleton. To demonstrate feasibility and evaluate system performance, we conducted an experiment in which one participant with a transtibial amputation tracked desired wrist trajectories during walking, while we measured wrist and ankle response.

**Results:** Benchtop testing demonstrated that for relevant walking frequencies, system error was below human perceptual error. During the walking experiment, the participant was able to voluntarily follow different wrist trajectories with an average RMS error of 1.55° after training. The ankle was also able to track desired trajectories below human perceptual error for both position control (RMSE = 0.8°) and torque control (RMSE = 8.4%).

**Conclusion:** We present a system that allows a user with amputation to control an ankle-foot prosthesis and receive feedback about its state using a wrist exoskeleton, with accuracy comparable to biological neuromotor control.

**Significance:** This bilateral teleoperation system enables novel prosthesis control and feedback strategies that could improve prosthesis control and aid motor learning.

## I. Introduction

MORE than 600,000 people live with major lower-limb amputation in the United States, a number that is expected to double by 2050 given the rising rates of vascular disease that lead to amputation [1]. As the primary cause of amputation in the US, vascular disease leads to reduced aerobic capacity and makes even slow walking a demanding task [2]. Those walking with conventional passive prosthetic limbs expend 20-47% more energy and have slower self-selected walking speeds compared to unimpaired individuals [3], [4]. Walking fatigue is second only to residual limb pain among concerns of those with lower limb amputation [5]. Limited mobility results in numerous secondary health problems and loss of independence, increasing medical costs and reliance on caregivers [6]. Additionally, people with amputation fall almost twice as much as those in the elderly population [7], [8].

Lower limb loss disrupts not only normal motor function, but also many sensory pathways. The human ankle, for example, is a complex joint comprised of muscle actuators and their attachments, in addition to sensory components, including muscle spindles and Golgi Tendon organs that relay information about the orientation and force production at the joint. Inputs to and from the central nervous system are also important, as the brain receives a copy of motor commands sent to the muscles to more accurately predict where the joint is in space [9]. An internal model then maps motor commands to expected sensory consequences [10].

Despite the complex interplay between sensorimotor commands in the biological ankle, most commercial ankle-foot prostheses focus primarily on restoring motor function to the user and lack sensory feedback from the joint. Recently, it has been shown that sensory feedback from a prosthestic limb can allow the user to experience more ownership of their limb [11], as well as reduce task times, metabolic cost, and phantom limb pain [12], [13]. The sensory feedback provided in these studies is typically either in the form of simplistic binary cues, such as vibrotactile [11] or electrocutaneous stimulation [12], or invasive surgical procedures [13], [14]. Because there is evidence that continuous feedback can result in reduced task times compared to binary feedback [15], and surgical options are expensive and invasive, there is room for development of new nonsurgical, continuous feedback methods for these devices. This type of feedback may be beneficial for the user long-term, and also allows us to design studies to learn more about sensory pathways of people with amputation. This type of feedback has been investigated for prosthetic hands by transmitting torque to a user’s elbow [16] or force to a user’s toes [17], but limited work has been done in the lower extremity.

In addition to the lack of sensory feedback provided by the majority of lower limb prostheses, most commercial devices are passive and therefore lack the ability to provide the net work or power that the biological ankle does during walking. Several powered ankle-foot prostheses exist, and one has been shown to reduce the metabolic cost of walking under some circumstances [18], [19], but how best to control them remains an open question. Usually these devices attempt to mimic typical behavior of an intact ankle during walking. There is reason to believe that customization could improve on this control, because studies using human-in-the-loop optimization in healthy individuals have shown that small individualized changes in kinematics or kinetics of an exoskeleton can result in large changes in metabolic cost [20]. However, pilot studies using a similar approach to optimize prosthesis parameters have resulted in negligible changes in metabolic cost [21]. Perhaps sensory feedback, with the addition of volitional control, is necessary for adaptation to these controllers.

Few studies have examined the benefits of providing the user with direct control over the movement of their lower limb prosthesis. Several surgical procedures, including targeted muscle reinnervation [22] and agonist-antagonist myoneural interfaces [23], show promise to improve control of prostheses in pilot studies, but these are invasive and expensive. The use of electromyography (EMG) from residual limb muscles as an input to the command signal for lower limb prostheses has also been tested [24]–[26]. However, measuring lower limb EMG requires placing sensors directly on muscles, many of which are loaded during walking because of their location within the socket of the limb. However, lower limb EMG requires placing sensors directly on muscles being loaded during walking, many of which are inside the prosthetic socket. This exacerbates signal disturbances such as changes in electrode position or loss of electrode-skin contact. The signal thereby degrades over time; existing systems must either be recalibrated or detect the signal degradation over time so they can revert to intent recognition through mechanical means [27], [28].

Teleoperation has been demonstrated as a highly effective way for people to directly control robotic devices when autonomy is not sufficient for the application [29]. Teleoperation allows for various combinations of force and position control pathways and feedback, and requirements for system stability are now well known [30], [31]. Studies have shown that teleoperation is effective in applications for upper-limb prostheses [32], in robot-assisted surgical systems [33], [34], and for rehabilitation, with information crossing between limbs [35]. Different modes of control are used in these applications, but all typically use rigid end-effectors with non-backdrivable actuation for both the manipulandum and the remote robot. Teleoperation of lower-limb exoskeletons has been investigated, but only in a virtual environment [36]. We propose to use a wrist exoskeleton to both teleoperate and receive sensory feedback about the state of a prosthetic ankle while walking (Figure 1). Such a system would allow us to answer scientific questions about sensory feedback and control for people with transtibial amputation and has the potential to improve user performance in terms of walking speed, balance, energy expenditure, and phantom limb pain.

**Fig. 1.**
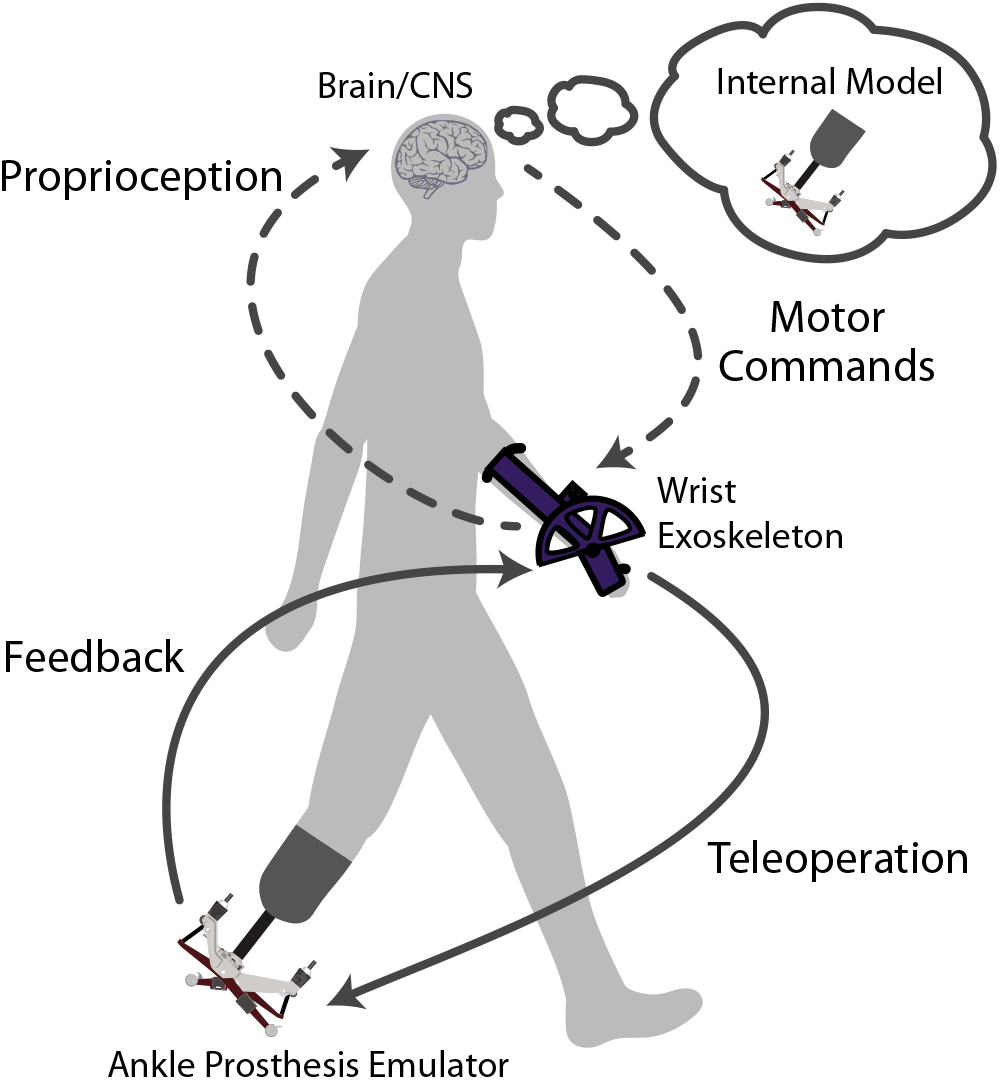
The wrist exoskeleton allows the user to control the ankle prosthesis, as well as receive augmented sensory feedback about the ankle prosthesis’ state, mimicking the control-feedback loop present in the unimpaired ankle. This augmented information being sent to the brain through the central nervous system (CNS) could allow the user to develop an internal model about the state of the ankle prosthesis and be able to better predict its motion.

The contributions of this work are: (1) the mechanical design of a wearable exoskeleton that is able to sense wrist angle and accurately apply wrist flexion and extension torques, (2) the development of control strategies for a novel teleoperation system that accounts for prosthesis actuator compliance and uncertainty in applied forces due to variations in the user’s gait, (3) benchtop tests characterizing the behavior of the wrist exoskeleton and ankle prosthesis, and (4) a feasibility study with a participant with amputation, quantifying the behavior of the system and the ability of the participant to voluntarily modulate ankle movements using the wrist exoskeleton during gait. Each contribution is addressed in further detail in the following sections.

## II. System Design

Our system consists of an ankle-foot prosthesis emulator powered by off-board motors, a wrist exoskeleton, and a computer to control both devices (Figure 2A). The ankle-foot prosthesis emulator, described in further detail in Section 2B, was previously designed and tested [37] (Figure 2B). In addition, we built a one-degree-of-freedom wrist exoskeleton capable of interfacing with the ankle-foot prosthesis emulator (Figure 2C-2D). Although there are many approaches that we could have taken to enable direct control, we chose the wrist for multiple reasons. First, the wrist joint in the arm is analogous to the ankle joint in the leg, and these joints have been shown to be linked in both interlimb reflexes [40] and brain activity [41]. Because of this neural coupling, we expected the wrist to facilitate more intuitive control than other upper-extremity joints. Controlling the prosthesis using the elbow or shoulder would likely disrupt arm swing, which is important for efficient gait [42]. Controlling the prosthesis using the hand or fingers could make the device less practical, and such mechanisms have proven difficult for participants to use to control exoskeletons [43]. With this in mind, we chose to design an exoskeleton controlled by wrist movement.

**Fig. 2.**
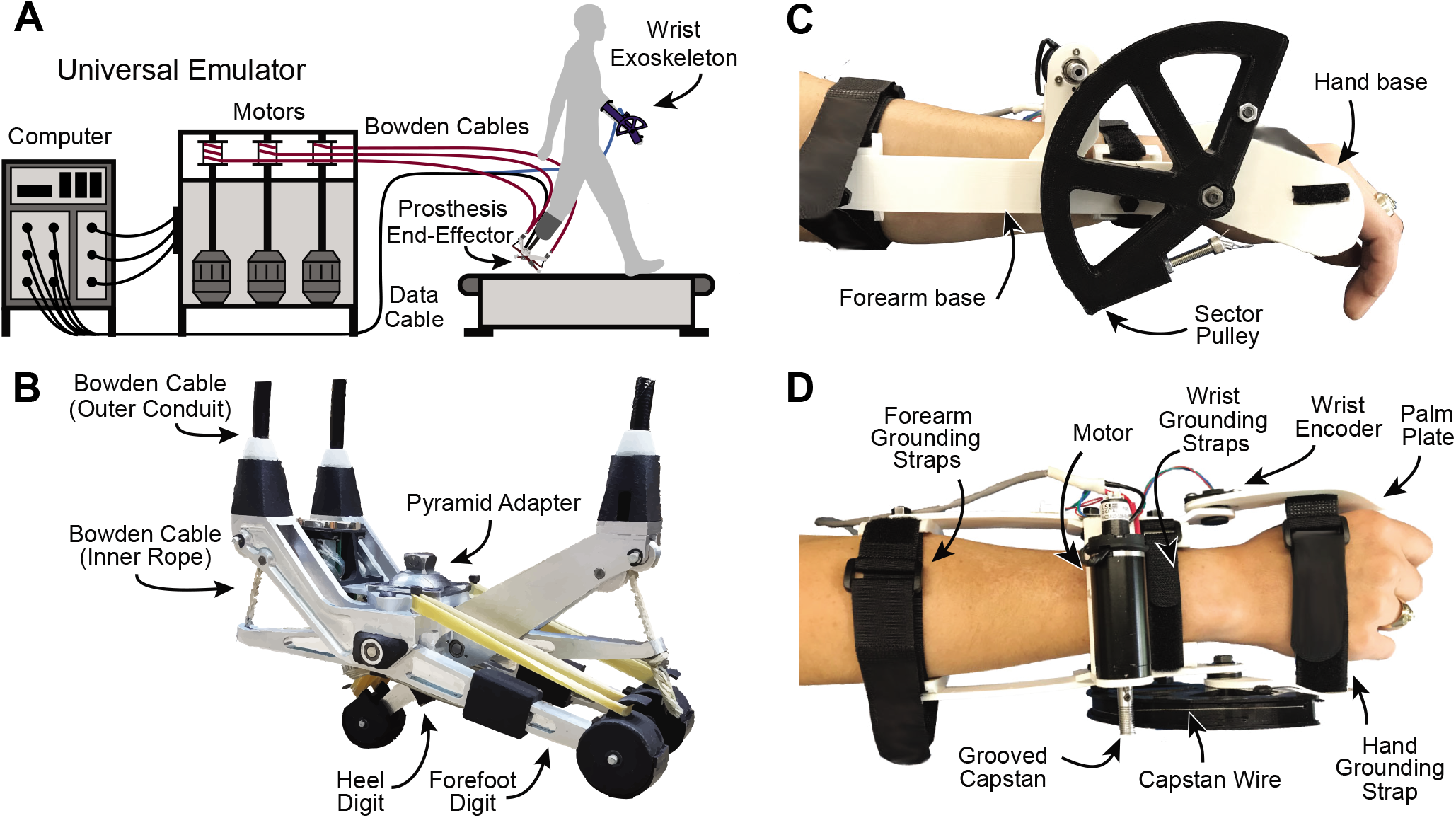
A. Schematic of the system containing the ankle-prosthesis emulator with off-board motors, the wrist exoskeleton, the user, and the computer that runs the controller. B. Position of the three digits of the ankle-foot prosthesis emulator is dictated by the tension from Bowden cables. C. Side view of the wrist exoskeleton shows that it is comprised of three separate, 3D-printed parts. D. Top view of the wrist exoskeleton shows the capstan motor drive, encoder, and grounding to the user’s arm.

### A. Exoskeleton Design

We had three primary design goals for the wrist exoskeleton, incorporating both the user interface and control fidelity. First, the wrist exoskeleton should be comfortable and lightweight to allow for natural motion of the arm. To achieve this goal, all base components were designed for mass efficiency and 3D printed from lightweight polylactic acid (PLA). The wrist exoskeleton comprises a forearm base and a hand base, with rigid links positioned on either side of the user’s arm (Figure 2C). Both are attached to the arm with Velcro straps, and the hand base is also grounded to the palm with a plate on the ventral side. The entire exoskeleton weighs 363 grams. To increase comfort and account for varying anthropometry of the forearm, spacers of different sizes can be attached to the inner portion of the wrist exoskeleton.

Our second design goal was to continuously transmit torques with an accuracy better than human wrist torque perception, while maintaining backdrivability. Device torques should be noticeable to the user, but significantly lower than the user’s maximum wrist flexion and extension torque, for both safety and to prevent fatigue. The average human is capable of 4.6 N·m of isometric wrist extension torque and 6N·m of isometric wrist flexion torque [38], so we chose 1 N·m as our target maximum torque. This allows the average user to overpower the exoskeleton by a factor of approximately five. In addition, human torque sensitivity at the wrist as a fraction of the applied torque has been shown to increase at higher torque magnitudes. Humans can detect a 12-13% change in the highest previously characterized reference torque of 0.3 N·m [39], so we use this as our target threshold for all torque magnitudes above 0.3 N·m. To achieve the desired torque output of 1 N·m design goal and maintain backdrivability, we used a capstan drive transmission. The capstan drive transmits torque via a flexible, inextensible cable from a grooved capstan, which is attached to the motor shaft, to a sector pulley (Figure 2D). The torque is amplified by the ratio of the capstan to the sector pulley radius. The exoskeleton is driven by an RE-25 motor (Maxon Motor, Switzerland) with a capstan ratio of 27 from the capstan pulley to the sector pulley diameter. To reduce interference with arm swing, we placed the capstan drive on the dorsal and lateral sides of the arm, which are furthest from the torso during natural arm swing. We also built a custom capstan pulley with grooved slots to minimize capstan wire slip.

The third design goal was to accurately measure wrist angle within the resolution of human proprioception, and to allow for full range of motion in wrist flexion and extension. Because the human wrist has three degrees of freedom and the wrist exoskeleton can only move in one degree of freedom, we inherently restrict wrist range of motion in radial/ulnar deviation and pronation/supination. The range of motion in wrist flexion and extension is a function of the sector pulley arc length. We chose an arc length that allows the user to achieve 75° of wrist flexion and extension, similar to a typical range of motion [40]. An RM08 encoder (RLS, Slovenia) on the joint opposite to the capstan drive measures wrist angle with a resolution of 0.18° over 180° of motion. Studies of human wrist proprioceptive resolution have reported values between 1.33° and 4.64° for flexion and extension [41]–[43], so the exoskeleton has significantly greater angle sensing resolution than humans.

### B. Ankle Prosthesis Emulator

This wrist exoskeleton interfaces with an ankle-foot prosthesis emulator previously described in [37] (Figure 2A-2B). The prosthesis emulator is a 3-DOF device with one heel and two forefoot digits, and a maximum plantarflexion and dorsiflexion angle of 19°. The device weighs 1.2 kg, and is capable of supplying 140 N·m of torque at the toes and 100 N·m of torque at the heel, using off-board motors (Kollmorgen Corp, Maryland, USA) that power the device via Bowden cables. This device is equipped with both an encoder and a strain gauge at each digit to measure angle and torque.

## III. System Control

There are several different strategies that could be used to control the ankle prosthesis with the wrist exoskeleton. Because we were unsure how well users would be able to successfully manipulate the ankle prosthesis if given full control, we developed two different control schemes. The first used direct position control with torque feedback, giving the user as much direct control and sensory information about the ankle prosthesis as possible but possibly making control more challenging. The second used torque control with a virtual spring, allowing the prosthesis to behave semi-autonomously, but enabling the user to alter its behavior with the wrist exoskeleton. We developed and tested the low-level controllers necessary to make these two control schemes possible. Figure 3 provides an overview of the controllers tested, and Table 1 provides a description for the symbols used in the following section. All control was done with a real-time target machine (Speedgoat, Switzerland) sampling at 1000 Hz.

**Fig. 3.**
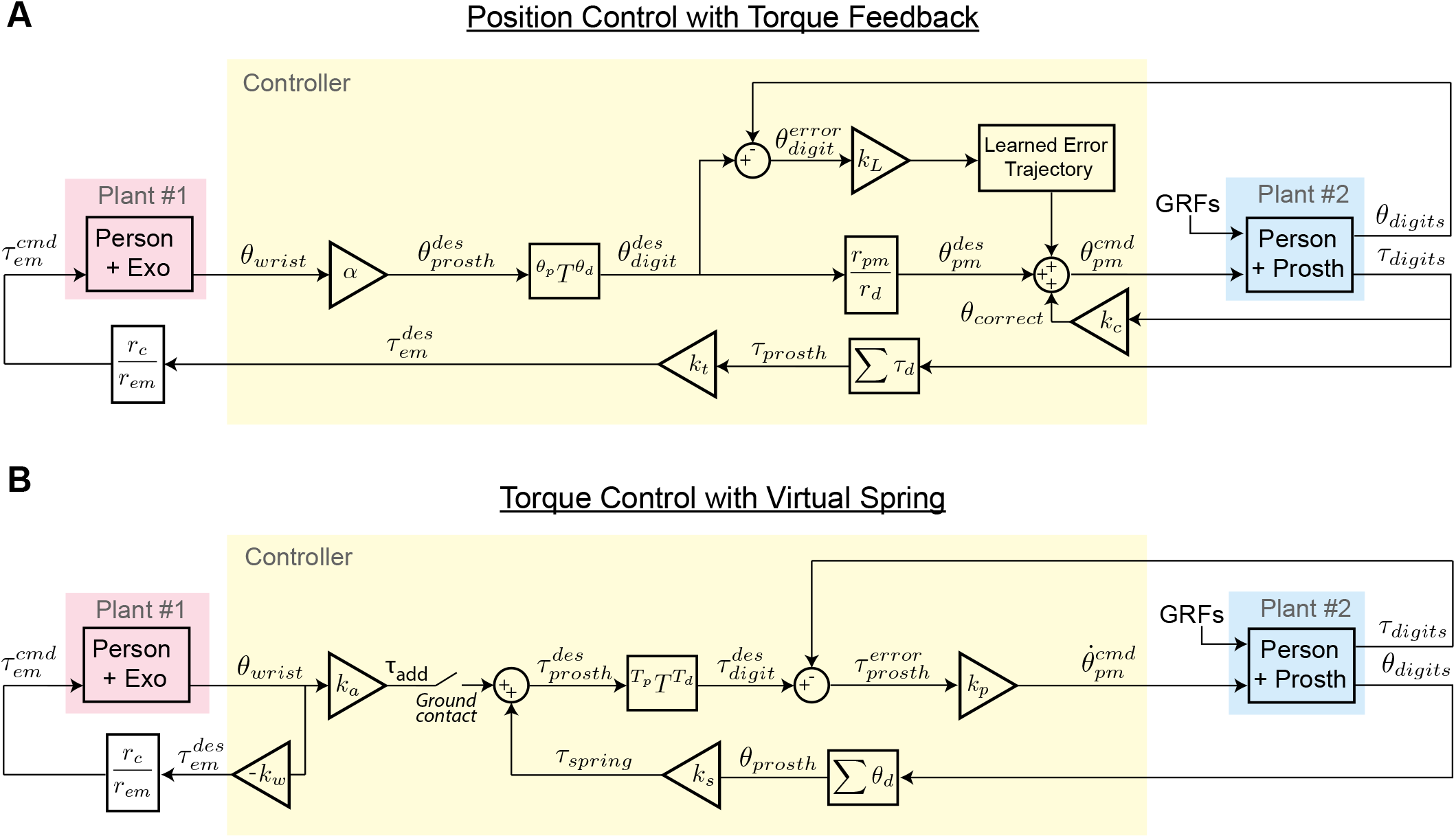
Control block diagrams of both high-level control schemes are shown. The diagrams are simplified to only show the control of one of the three ankle prosthesis digits. A. In position control, the user directly controls the position of the prosthesis (Prosth) with the wrist exoskeleton (Exo). This is accomplished by transforming a desired prosthesis angle to a desired position of the motor drum that controls the Bowden cables, using model-based and model-free corrections. When haptic feedback is provided, a scaled version of the ankle prosthesis torque is fed back to the wrist exoskeleton for both controller types. B. In torque control, the user exerts force against a virtual spring in the wrist exoskeleton, which is transformed to an ankle torque added to a basic spring controller.

**Table 1:**
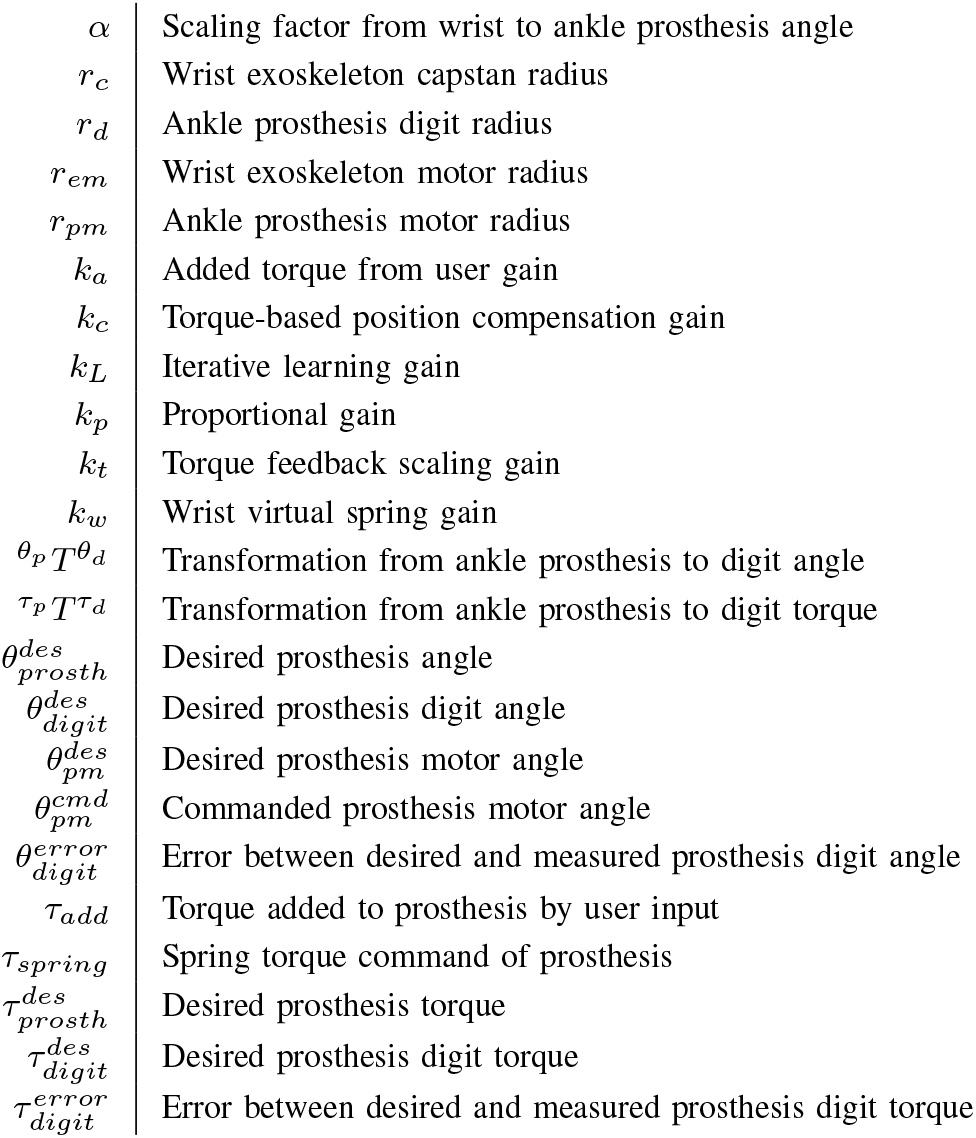
Variables and Parameters

|  |  |
| --- | --- |
| $\alpha$ | Scaling factor from wrist to ankle prosthesis angle |
| $r_c$ | Wrist exoskeleton capstan radius |
| $r_d$ | Ankle prosthesis digit radius |
| $r_{em}$ | Wrist exoskeleton motor radius |
| $r_{pm}$ | Ankle prosthesis motor radius |
| $k_a$ | Added torque from user gain |
| $k_c$ | Torque-based position compensation gain |
| $k_L$ | Iterative learning gain |
| $k_p$ | Proportional gain |
| $k_t$ | Torque feedback scaling gain |
| $k_w$ | Wrist virtual spring gain |
| $\theta_p T^{\theta_d}$ | Transformation from ankle prosthesis to digit angle |
| $\tau_p T^{\tau_d}$ | Transformation from ankle prosthesis to digit torque |
| $\theta_{prosth}^{des}$ | Desired prosthesis angle |
| $\theta_{digit}^{des}$ | Desired prosthesis digit angle |
| $\theta_{pm}^{des}$ | Desired prosthesis motor angle |
| $\theta_{pm}^{cmd}$ | Commanded prosthesis motor angle |
| $\theta_{digit}^{error}$ | Error between desired and measured prosthesis digit angle |
| $\tau_{add}$ | Torque added to prosthesis by user input |
| $\tau_{spring}$ | Spring torque command of prosthesis |
| $\tau_{prosth}^{des}$ | Desired prosthesis torque |
| $\tau_{digit}^{des}$ | Desired prosthesis digit torque |
| $\tau_{digit}^{error}$ | Error between desired and measured prosthesis digit torque |

### A. Position Control with Torque Feedback

In this control scheme, the user controls the position of the ankle using the wrist to provide the reference input, while simultaneously receiving torque feedback from the ankle. Therefore, the user receives proprioceptive feedback about the angular position of the ankle via their wrist proprioception, in addition to feedback at the wrist regarding the ankle torque.

The ankle prosthesis has three degrees of freedom (the heel and two toes), while the user only commands one degree of freedom, so the user’s wrist angle is mapped to a single commanded ankle angle. The wrist position command is first converted to a commanded ankle angle by multiplying by a scaling factor, *α*, because the typical range of motion of the wrist is much larger than the range of motion of the ankle. Wrist extension corresponds to ankle dorsiflexion, and wrist flexion corresponds to ankle plantarflexion, as shown in Figure 4A. This commanded ankle angle is then translated to angular positions for each of the digits, using two additional constraints on the position: (1) a set offset between the two toe digits, and (2) a set overall height for the prosthesis, approximately equal to the height of the intact ankle joint of the user with amputation. Both the toe offset and the fixed height are determined with the help of a prosthetist during an initial evaluation session.

**Fig. 4.**
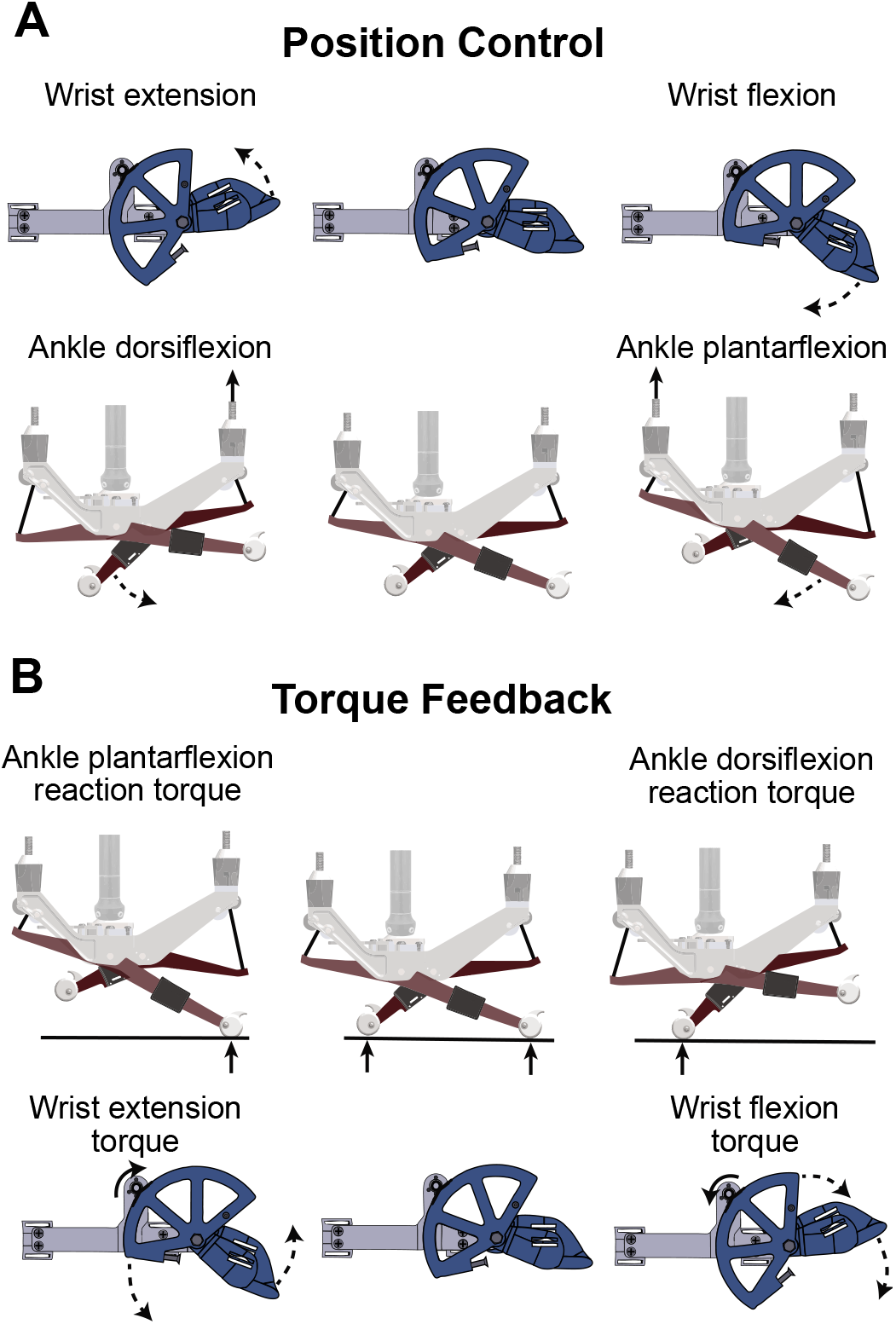
The mechanisms of position control and torque feedback are demonstrated above, with forces and torques displayed in solid lines and resulting changes in position in dashed lines. A. In position control, wrist extension results in greater ankle dorsiflexion by reeling in the Bowden cable connected to the heel of the prosthesis, resulting in downward motion of the heel. Wrist flexion results in greater ankle plantarflexion by reeling in the cables connected to the front toes of the prosthesis, resulting in downward motion of the toes. B. Ankle plantarflexion reaction torque produces greater forces on the toes of the prosthesis. To produce the torque feedback, the ankle plantarflexion reaction torque results in the wrist exoskeleton motor being driven clockwise to produce a wrist extension torque, generating an upward motion at the palm plate. In contrast, the ankle dorsiflexion reaction torque results in the wrist exoskeleton motor being driven counterclockwise, generating a downward motion at the palm plate.

The desired angle for each digit of the prosthesis is then computed and translated into a desired position of the motor drum, 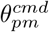 which dictates the length of the Bowden cable controlling the prosthesis digit. Using the relationship between the radius of the prosthesis digit and the radius of the motor drum, along with the initial voltage commanded at a starting position, we determine the input voltage required to reach a desired position. The effect of elasticity of the Bowden cables and forces applied at the digits as the user walks on the prosthesis is compensated for using two correction terms: model-based and model-free. The model-based term treats each Bowden cable as a simple spring, resulting in a linear relationship between forces applied at the digits and position errors. Therefore, the correction term of *k_c_τ_digit_* is added to the desired motor position. Because the simple linear model does not capture all errors, a second term provides model-free correction based on iterative learning. This additional learning term keeps a running average of the errors (*e*) accumulated at each timepoint in the gait cycle, which are used to apply a correction at each of those timepoints plus a pre-determined time delay throughout the gait cycle, multiplied by a learning gain, *k_L_*. This iterative learning approach has been previously described and implemented in cyclic walking tasks [20], [37]. The following control equation is used for the position control of the ankle prosthesis at each digit:

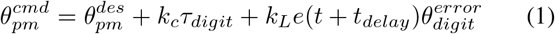

The wrist exoskeleton motor (em) can also receive scaled torque feedback from the ankle. The reaction torque resulting from forces on the toes in ankle plantarflexion is translated to a wrist extension torque, while the reaction torque resulting from greater forces on the heel is translated to a wrist flexion torque, as shown in Figure 4B. To transmit this torque, we use a simple proportional gain with a scaling factor of *k_t_*, to account for large torques at the ankle that would be unsafe and uncomfortable to transmit to the wrist:

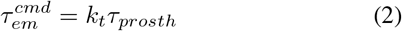

Because similar high-quality haptic devices have been shown to be effective in open-loop control [44], we use open-loop control for this feedback system following benchtop testing to ensure torque display accuracy similar to human torque perception accuracy.

### B. Torque Control with Virtual Spring

In this control scheme, the ankle prosthesis tracks a simple spring controller, while the user has the ability to modulate ankle prosthesis torque using the wrist. The following control law dictates the behavior of the ankle prosthesis:

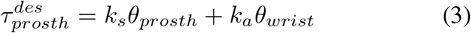

The *k_s_* gain determines the stiffness of the spring that governs the baseline motion of the ankle prosthesis, while the *k_a_* gain determines the magnitude of the additional torque added or subtracted by the motion of the user.

The low-level control of each digit of the prosthesis consists of a proportional feedback term in velocity control. This is governed by the following equation:

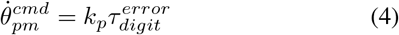

The haptic feedback provided in this control mode is a virtual spring implemented at the wrist, which provides increasing torque to the wrist the further the wrist is driven away from the zero position. This allows the user to feel a scaled version of the torque that they are adding or subtracting from the device, and demonstrates where the neutral (zero) position of the wrist exoskeleton lies. This virtual spring is governed by the following equation:

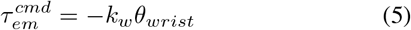

## IV. Benchtop Testing

We performed benchtop testing of the behavior of both the wrist exoskeleton and ankle prosthesis when controlled as described in Section III. Torque tracking accuracy of the wrist exoskeleton was tested by comparing input signals to known torque outputs. In addition, for each device and control mode, a frequency response test was performed to determine how the system behaves across various input frequencies. Novel characterization tests were performed for the position response of the ankle prosthesis emulator and the torque response of the wrist exoskeleton. The torque response of the ankle prosthesis emulator has been previously characterized [37], but we present it here as well for comparison.

Our target goals for control accuracy were as follows: (1) static accuracy within the threshold for human perception, and (2) dynamic accuracy within the threshold of human perception for input frequencies under 6 Hz. We chose 6 Hz because the majority of the frequency content is below this threshold during walking [45]. Because of this design goal, we present a sample trace of the input and output values at this frequency (Figure 5A), in addition to dynamic accuracy plots across all tested frequencies (Figure 5B). To set target thresholds, we used human perception data from literature. Ankle proprioceptive errors have been reported to be 2.3° [46]. Human error in torque perception is typically characterized as a just noticeable difference (JND) that varies depending on the applied reference torque. For wrist flexion and extension torque with reference magnitudes similar to those used in benchtop testing, a JND value of 0.04 N·m has been reported [39]. To our knowledge, no one has directly examined the JND for ankle plantarflexion and dorsiflexion torque. However, it has been shown that humans can reliably detect stiffness changes of greater than 12% at the ankle [47], so we use 12% of the maximum applied torque as our target for ankle torque perception.

**Fig. 5.**
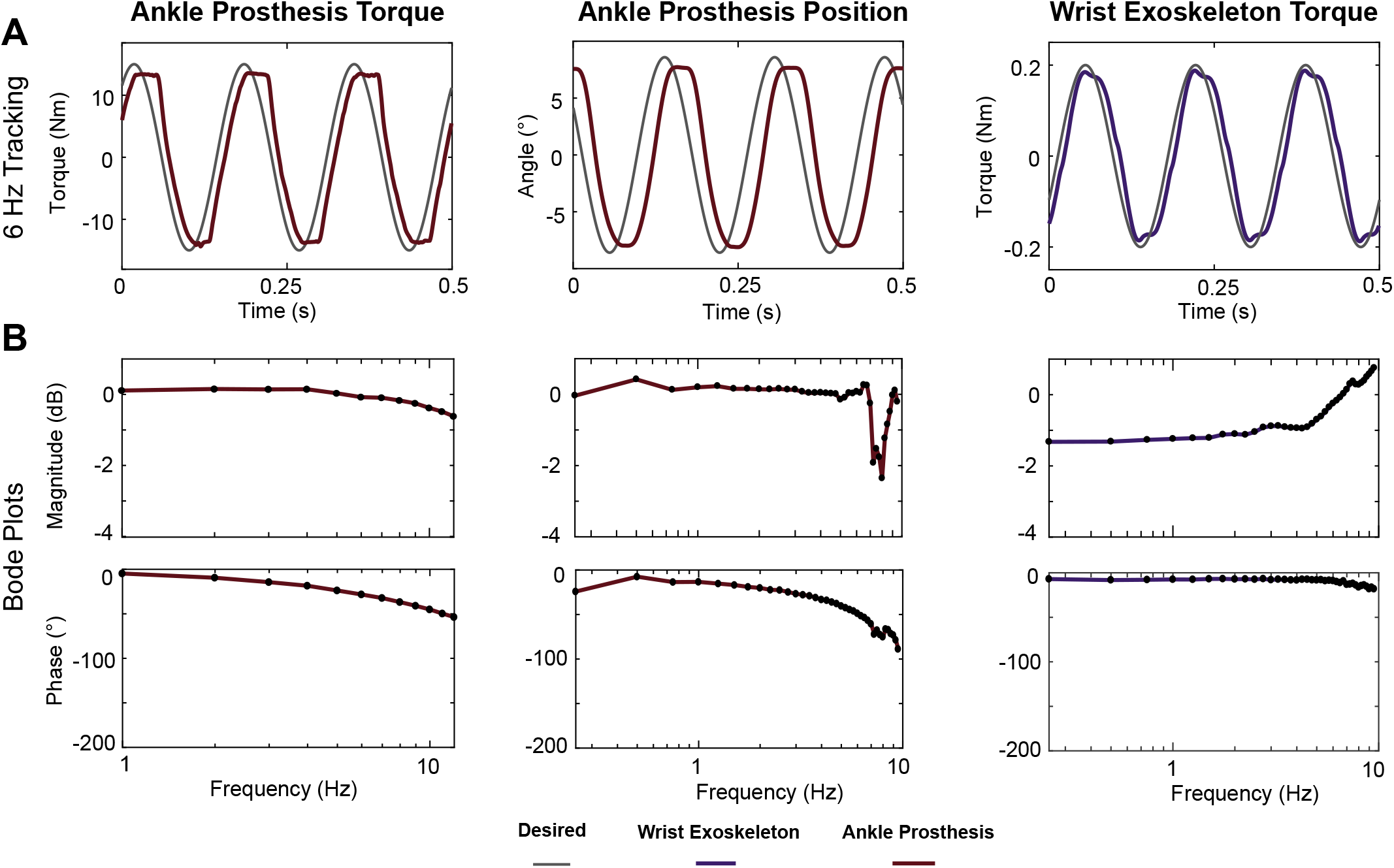
Results from benchtop testing of both the ankle prosthesis and the wrist exoskeleton in each control mode: ankle prosthesis torque, ankle prosthesis position, and wrist exoskeleton torque. A. The majority of the frequency content in walking is below 6 Hz, and a sample trace at this frequency shows that the output signals track the desired curve with reasonable accuracy. B. Bode plots demonstrate that the bandwidth of each system is greater than 6 Hz.

Input frequencies up to 10 Hz were tested in increments of 0.25 Hz, except for the previously characterized ankle prosthesis torque, which was tested in increments of 1 Hz. Each frequency was commanded for 3 seconds, and the output was fit to a sine wave. The resulting amplitude and phase shift were used to generate a Bode plot. We define bandwidth as the lowest frequency during which the amplitude ratio drops below −3 dB or the phase margin exceeds 150°. In order to provide the most conservative estimate of performance, iterative learning was not used during benchtop testing.

### A. Torque response of ankle prosthesis emulator

For torque response testing of the ankle prosthesis emulator, the end-effector was fixed in a rigid frame to prevent movement, as described in [37]. Although the testing for each digit was performed separately, the responses of all digits were identical, and therefore only one result is shown for each test. Measurement error was evaluated by comparing known applied torque to torque measured by the prosthesis emulator using strain gauges. Root-mean-square (RMS) measurement error was 1.7 N·m. Because the maximum torque applied during benchtop testing was 15 N·m, this resulted in an error of 11.3%, less than our target of 12%. Up to 10 Hz, the magnitude and frequency response of the system had high fidelity, with the magnitude response degrading by less than 1 decibel and the phase lagging by less then 50° (Figure 5B). The response to a 6 Hz input is shown in Figure 5A.

### B. Position response of ankle prosthesis emulator

For all position response characterization tests, the prosthesis was fixed in midair so that all digits could move freely. As described in Section III, a position input combined with a set height was used to command all three digits simultaneously to result in an overall ankle angle proportional to the input. The proportional and derivative gains were held constant during all tests. Based on these calculations, we determined the position control bandwidth to be greater than 10 Hz, which exceeds our target of 6 Hz (Figure 5B). The response to a 6 Hz input is shown in Figure 5A. To find the position sensing accuracy of the ankle prosthesis, we used the accuracy of the RM08 encoders on each digit, which have a resolution of 0.18°, less than human proprioceptive error of 2.3°.

### C. Torque response of wrist exoskeleton

To measure the accuracy of the motor torque applied to the wrist exoskeleton, we commanded a virtual spring centered around a neutral angle and hung masses of known values from the wrist base, such that the further the motor traveled from the neutral position, the more resistance torque was applied. Each mass was allowed to reach steady state, and the motor torque commanded was averaged. This averaged torque was compared to the known torque resulting from the mass hanging on the motor shaft with a known radius, compensating for the change in angle as a result of the displacement of the wrist base. This test was repeated in triplicate with 5 known masses. Both the wrist flexion and extension torque were tested by fixing the wrist exoskeleton upside-down in order to test the opposing direction. The resulting fit for the torque values was very linear, with an *R*^2^ value of 0.992. In addition, the RMS error between commanded and actual torque was 0.0305 N·m, which is less than our target value of 0.04 N·m.

The open-loop torque frequency response test was conducted using an external 6-axis Nano17 force/torque sensor (ATI Industrial Automation, North Carolina, USA). To conduct this test, a separate wrist exoskeleton was built identical to the original, but which housed a force/torque sensor instead of the encoder in the opposite joint to the capstan drive. The main frame of the wrist exoskeleton was grounded in order to minimize movement during testing. In the range of our testing frequencies, we did not reach the bandwidth of the system (Figure 5B). The Bode plot revealed an increasing magnitude response while the phase decreases. Based on the behavior of other haptic devices with capstan drive mechanisms, we expect that the increasing magnitude is a result of approaching the resonant frequency of the device, after which the magnitude would decrease. The response to a 6 Hz input is shown in Figure 5A.

## V. Walking Trial

We recruited one participant with a left-foot transtibial amputation (male, 44 years old, 2 years post-amputation) to walk with the devices on a treadmill (Bertec, Ohio, USA). The participant used the wrist exoskeleton on his right wrist to control the ankle-foot prosthesis on the contralateral leg in both position control and torque control. In this pilot experiment, we were interested in (1) the accuracy with which our system was able to control desired ankle angle or torque during walking, and (2) the accuracy with which the participant was able to command desired wrist angle. It has been previously demonstrated that humans can use real-time visual feedback to modulate their gait patterns [48], [49] and upper extremity movement [50]. However, studies instructing subjects to modulate their gait typically provide cues in the form of binary feedback, and studies in the upper extremity typically occur while participants are seated. Therefore, we measured how well the participant could follow specific continuous wrist trajectories in real-time while walking. Prior to testing, the ankle-foot prosthesis was fit to the participant by a licensed prosthetist. All tests were done following a protocol approved by the Institutional Review Board at Stanford University, and the participant gave written informed consent.

### A. Experimental Protocol

Training and testing for the experiment was completed over the course of two days. On the first day of the experiment, the participant acclimated to the system, then we tested torque control. The position controller was tested on the second day. For each type of control, the participant completed multiple training trials to practice teleoperating the ankle while seated or standing before walking. In addition, for the position control condition, he first completed training and testing trials without haptic feedback before haptic feedback was added, both to allow the user to acclimate to the control first, in addition to comparing the system behavior with and without the feedback. During each training or testing block, the participant completed two trials of five minutes each, following two separate wrist trajectories, explained in further detail in Section VB. An overview of the training and testing completed for each type of control is shown in Figure 6. During all walking trials, the participant was allowed to self-select his walking speed, which was between 0.8 m/s and 1.0 m/s across both days.

**Fig. 6.**
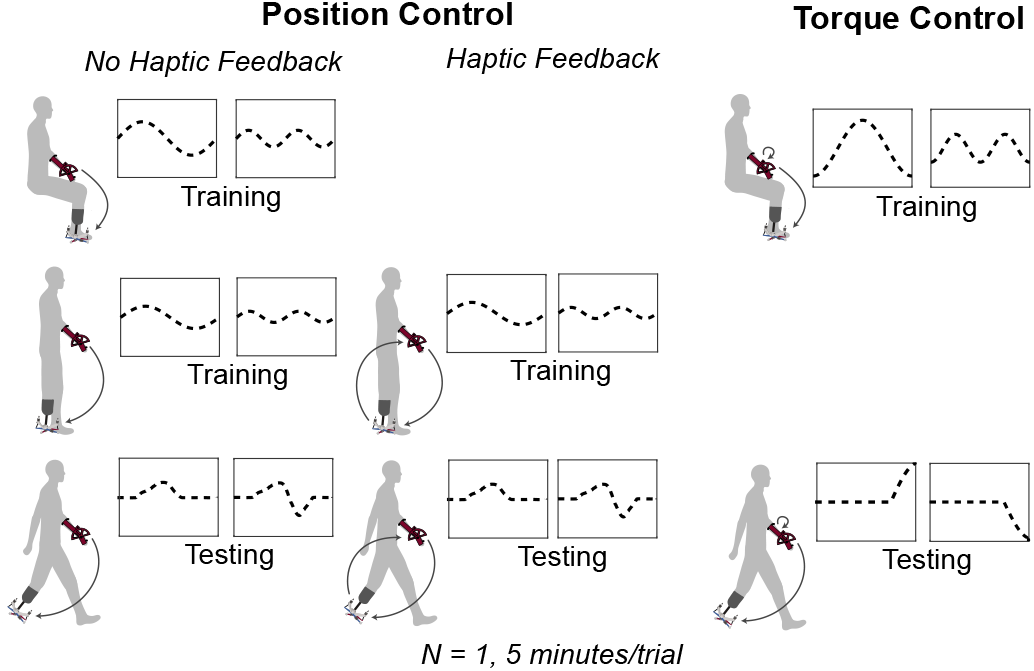
The training and testing protocol for each type of control is shown. Training trials allowed the participant to practice each type of control while seated or standing before walking. All trials lasted 5 minutes, and two different trajectories were provided for each training or testing condition.

Iterative learning was turned on only in the last 90 seconds of the position control walking trials. We allowed the participant to first walk without iterative learning in order to establish consistent cyclic errors, and found that 3.5 minutes allowed the participant to achieve a consistent desired wrist, and thus ankle, trajectory. This allowed for effective error compensation using iterative learning. All trials were successfully completed for the full 5-minute duration except for the position control trial with active push-off and no haptic feedback. This trial produced spikes in the torque profile and was ended 30 seconds early due to subject discomfort.

### B. Real-Time Feedback

In order to demonstrate that the participant was able to command different ankle trajectories with his wrist, we provided two different trajectories for him to follow in each training and testing condition. In all conditions, the user was able to see the desired wrist trajectory and the real-time wrist angle displayed on a 40-inch screen placed in front of him. We chose to display the desired wrist trajectory instead of the desired prosthesis trajectory in order to separate the human error between desired and realized wrist motion from the error in the mechanical system (between desired ankle prosthesis angle or torque and realized angle or torque). For each trial, we measured the root mean square error between the desired and measured wrist trajectory.

The desired wrist trajectories given for the training conditions were different from the test conditions, both in pattern and mechanics of how they were displayed. In all training trials where the subject was seated or standing, the desired trajectories were sine waves of various amplitudes and frequencies. The horizontal axis of the displayed graph was based on time, and so the real-time feedback to the subject about the current state of the wrist was reset after a set time period. The desired trajectories for the walking trials were based on previously published kinematic data from people without amputation [51]. For the position control conditions, we chose one trajectory emulating passive walking and another emulating active push-off. The horizontal axis used for position control was percent gait cycle, and the graph reset once a new heel strike was detected. For torque control, the user only had control of the ankle during the stance phase, so percent stance was used as the horizontal axis in the real-time feedback plot. Similarly to position control, two trajecories were chosen with differing amounts of ankle plantarflexion torque during push-off: one in which the user removed plantarflexion torque during push-off, and one in which the user injected additional platarflexion torque during push-off.

### C. Analysis

For both human wrist error and system ankle-prosthesis error, we were interested in the root mean square (RMS) error for each gait cycle in the last 30 seconds of each trial. In order to obtain an equal number of gait cycles for comparison between trials, we identified the trial that contained the fewest number of gait cycles in the last 30 seconds, and only included the average RMS error for this number of gait cycles for the other trials as well. For each trial, we performed a onesided t-test comparing the RMS errors from the end of the trial to the average human proprioceptive or kinesthetic error taken from literature, as described in Section IV for human wrist proprioceptive error and human ankle torque perception. In addition, although ankle proprioception is not directly comparable because the participant is not sensing ankle angle, we use the average human ankle proprioceptive error as a comparison to provide a benchmark for our ankle error in position control.

### D. Results

#### 1) Human Wrist Control

The average and standard deviation of RMS error in wrist position for the end of each testing trial is shown in Figure 7A. During the torque control condition, which was tested on the first day, wrist RMS error was greater than wrist proprioceptive error for both trials. However, average RMS errors during all position control trials tested on the second day, both without haptic feedback (Pos) and with haptic feedback (PosH), were significantly less than human wrist proprioceptive error (Pos Less Plantarflexion: *p* = 1.18 × 10^−8^, Pos More Plantarflexion: *p* = 6.04 × 10^−9^; PosH Less Plantarflexion: *p* = 1.25 × 10^−14^, PosH More Plantarflexion: *p* = 4.10 × 10^−14^). We hypothesize that the discrepancy between the two types of control is due to the subject having additional training with the system by the second day, instead of some inherent difference between the two types of control or trajectories provided. In addition to the grouped data, individual and averaged wrist angle traces for all gait cycles at the end of each trial are shown for position control (Figure 8) and torque control (Figure 9).

**Fig. 7.**
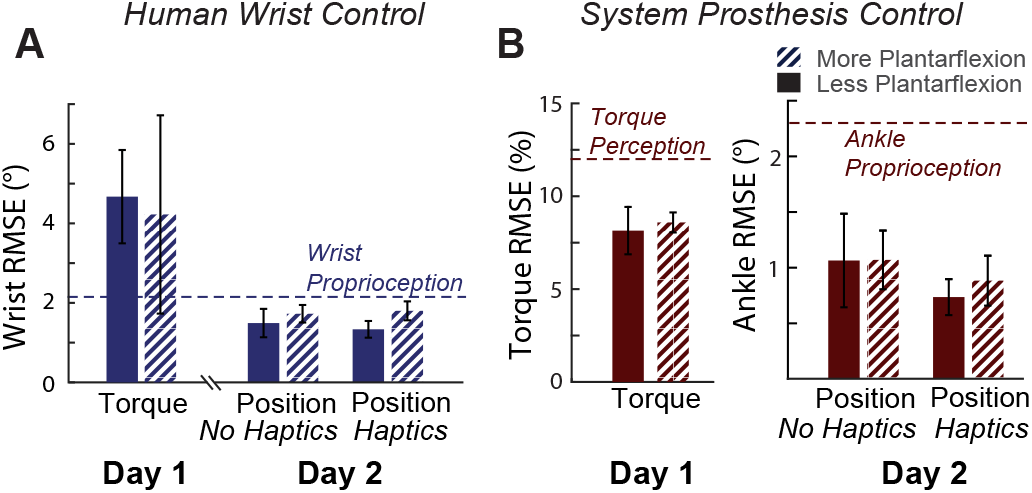
Average and standard deviation of RMS error for wrist position (A) and ankle prosthesis state (B) are shown. These are compared with human wrist proprioceptive or kinesthetic errors from literature. A. When torque control was tested on Day 1, wrist RMS errors were higher than wrist proprioceptive errors. However, by Day 2, when position control was tested, RMS error both without haptic feedback and with haptic feedback was significantly less than wrist proprioceptive error. B. Ankle prosthesis position error and ankle prosthesis percent torque error was significantly less than human perceptive errors during all torque control and position control trials.

**Fig. 8.**
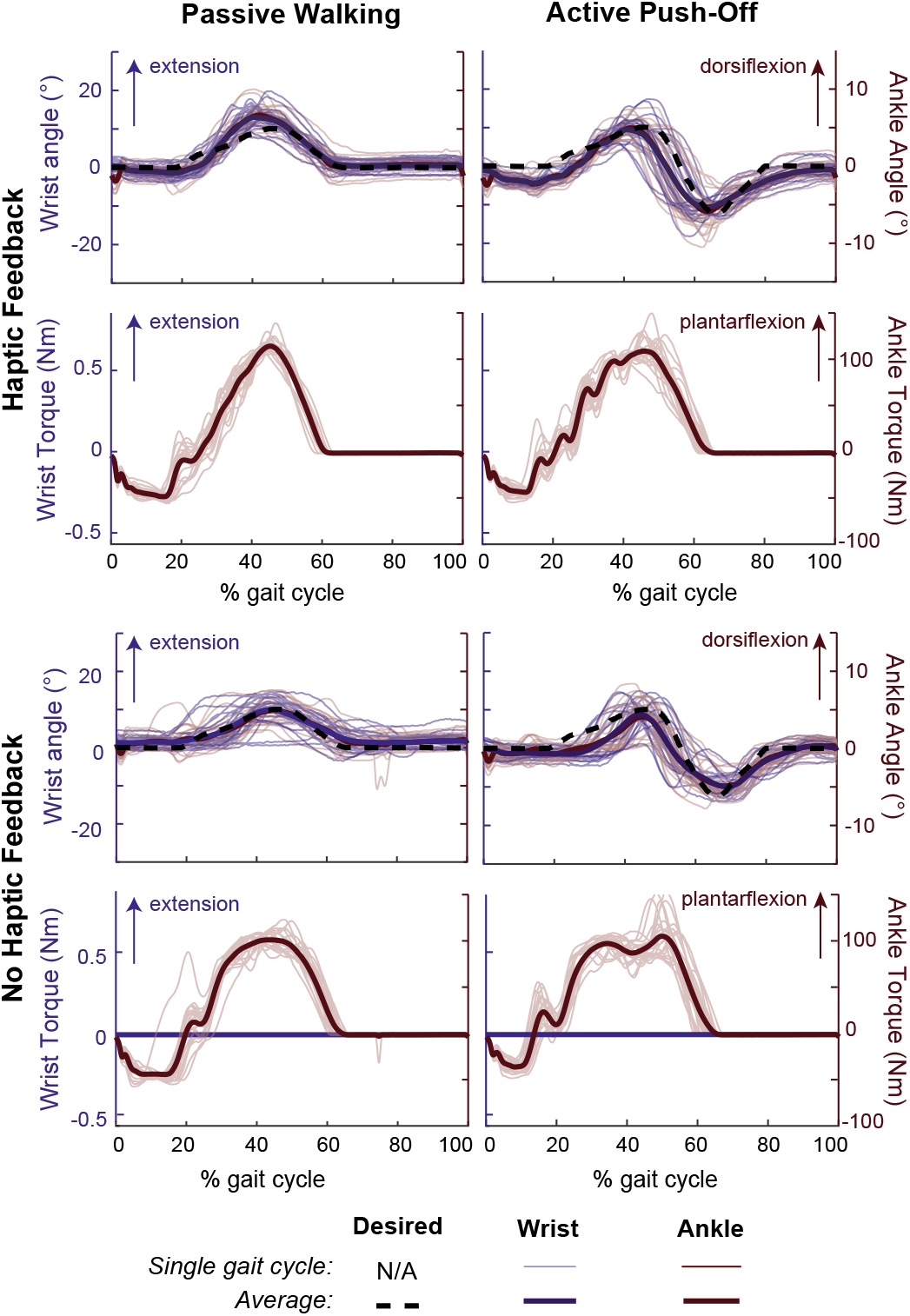
Angle and torque of the wrist exoskeleton and ankle prosthesis for each position control condition. These representative data are from the final 30 seconds of trials in which the target trajectory was most similar to biological gait (the same trials as presented in Figure 6). For each condition, desired wrist trajectory, wrist position, and ankle position are shown in the top plot, and ankle torque and commanded wrist torque for the feedforward torque control are shown in the bottom plot. Because we used the previously characterized wrist exoskeleton properties to estimate wrist torque, the commanded wrist torque in the haptic feedback conditions exactly matches the scaled version of the ankle torque, and for this reason the traces of the wrist torque are not visible.

**Fig. 9.**
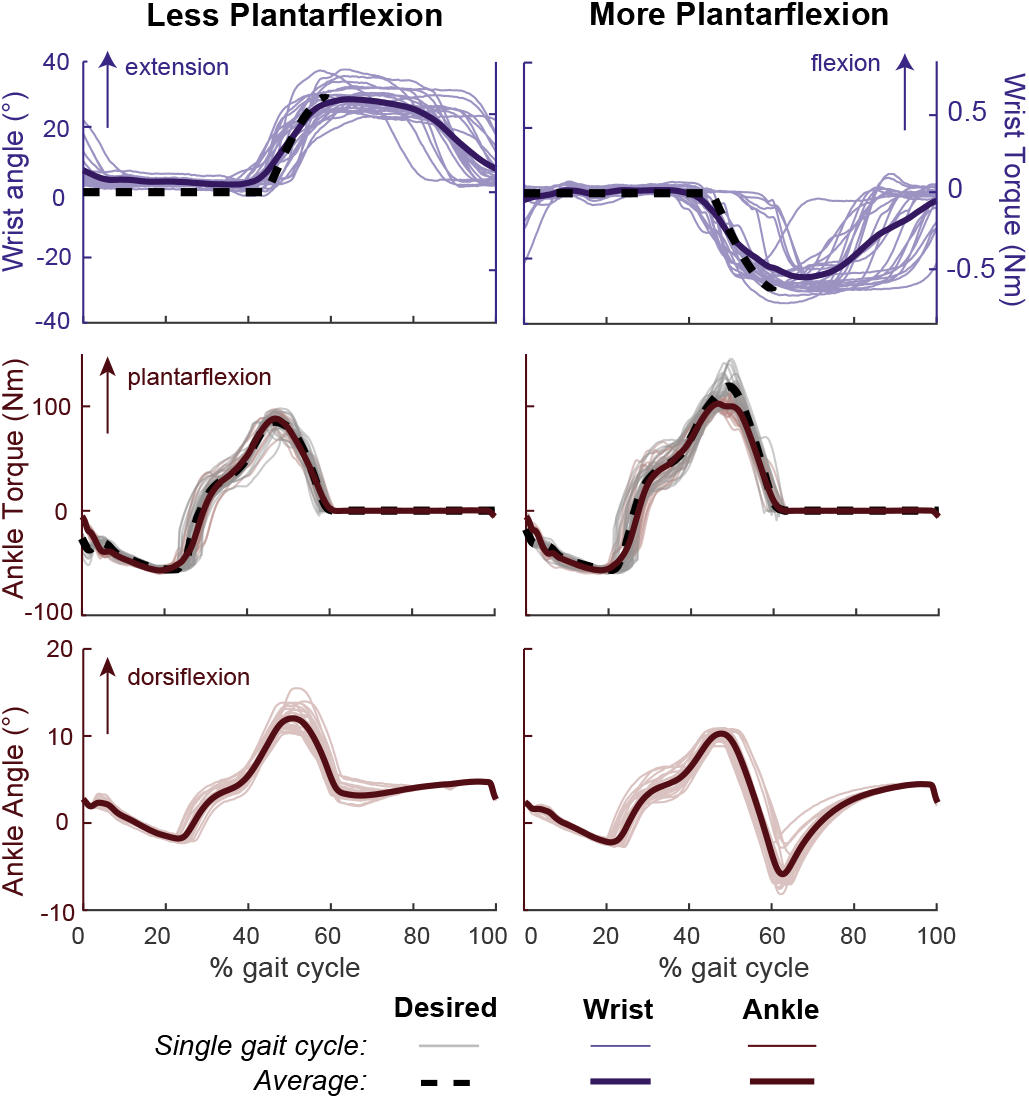
Data from the torque control condition is shown, with the trial corresponding to less plantarflexion on the left and the trial corresponding to more plantarflexion on the right. The top row shows the desired and realized wrist trajectories, the middle row shows the commanded and measured torque from the prosthesis, and the bottom row shows the ankle position. The two trajectories produced different torque trajectories, although there was some error between desired and measured torque.

#### 2) Prosthesis Position Control

As shown in Figure 7B, RMS error between commanded and realized ankle angle in all position control trials was significantly less than human ankle propriocpetive error (Position Control with No Haptics and Less Plantarflexion: *p* = 2.72 × 10^−12^, Position Control with No Haptics and More Plantarflexion: *p* = 3.06 × 10^−16^; Position Control with Haptics and Less Plantarflexion: *p* = 9.58 × 10^−23^, Position Control with Haptics and More Plantarflexion: p = 6.53 × 10^−19^). Figure 8 shows that although the commanded ankle angles followed similar trajectories for each trial in the haptic feedback and no haptic feedback conditions, the resulting ankle torques were qualitatively quite different. In addition, small oscillations are seen in the ankle torque profiles, particularly with the active push-off trajectory. Future work will investigate the cause of these oscillations to mitigate them.

#### 3) Prosthesis Torque Control

For the torque control trials, Figure 8 shows the desired and measured wrist trajectories, in addition to the resulting ankle torque and position. These data are shown for both trials, with more or less plantarflexion torque. As expected, when the participant commands more plantarflexion torque during push-off, the ankle angle plantarflexes and the average maximum torque increases, from 88.3 N·m to 102.5 N·m, showing that the participant was able to alter the torque trajectory of the ankle. Error in ankle torque tracking results in an ankle torque that is less than commanded at push-off for the trial commanding greater plantarflexion torque. The average RMS error between commanded and measured ankle torque as a percentage of the maximum ankle torque is 8.15% for the trial with less plantarflexion and 8.59% for the trial with more plantarflexion. As shown in Figure 7B, both of these values are significantly less than the human ankle error threshold for stiffness perception of 12% [47] (Less Plantarflexion: *p* = 2.18 × 10^−64^, More Plantarflexion: *p* = 3.09 × 10^−72^).

## VI. Discussion and Future Work

We developed a system that allows a user with a transtibial amputation to teleoperate their ankle-foot prosthesis and receive haptic feedback about the state of the prosthesis. A wrist exoskeleton senses wrist angle and implements wrist torque up to 1 Nm. Two different teleoperation schemes allow the wrist exoskeleton to interface with the ankle prosthesis. The first directly controls the ankle prosthesis angle and receives scaled wrist torques from the prosthesis. The second modifies a spring-like torque trajectory with the wrist and receives haptic feedback proportional to the torque that the user inputs or removes from the system. A person with a transtibial amputation was able to effectively use the wrist exoskeleton to teleoperate the ankle prosthesis in real time using these control schemes.

Of the two control schemes tested, the position control provides the user with more information because they are able to feel a scaled version of the ground reaction torque from the prosthesis at their wrist, in addition to using their intact wrist proprioception to estimate ankle angle. However, because the ankle prosthesis follows a scaled version of the wrist angle, the wrist movement needed to generate an ankle trajectory similar to the biological ankle is complex and therefore may result in greater cognitive load for the user. In contrast, the torque control scheme does not provide the user with as much information. The ankle prosthesis has a baseline behavior of a passive spring, and the user can inject or remove torque from this behavior via wrist movement. Because of the virtual spring at the wrist, the user can feel a scaled version of the torque that they are injecting or removing, but does not have a concrete representation of the overall torque or ankle position at any instant in time. While the user does not have as much information, the wrist trajectories required to generate a natural ankle trajectory can be much simpler. In future work, functional gait metrics should be measured with the control approaches we have developed, as well as haptic feedback alone, to examine their individual effects. In addition, the differences between cognitive load or comfort of different control schemes could be tested.

In our teleoperation control schemes, we control the behavior of two separate devices: the wrist exoskeleton and the ankle prosthesis. Yet because both devices are attached to the human user, the system actually has two plants that are each a combination of the device and the limb to which they are attached: (1) the wrist exoskeleton and the wrist, including all of its sensorimotor inputs and outputs, and (2) the ankle-foot prosthesis and sensorimotor inputs and outputs from the residual limb and rest of the body that affect gait and therefore ground reaction forces. Accurate control of the prosthesis depends not only on the mechatronic system capabilities, but also on the capability of the user to accurately control their wrist in real time while they are walking. We found that, by the second day of training, our participant was able to match multiple desired trajectories with errors less than that of human wrist proprioceptive errors. Because this was a proof-of-concept study with one participant, further work is required to generalize these results and characterize human adaptation to the system.

We were able to achieve sufficient position control accuracy with this system, with ankle position RMS errors less than human ankle proprioceptive errors. However, with this control strategy we noticed small oscillations in resulting ankle torque, especially with haptic feedback present. Other teleoperation systems have noted a trade-off between higher tracking accuracy and this type of oscillatory behavior [33]. Future work will examine this possible trade-off between position control accuracy and torque oscillations. Additionally, it is unclear if perfect position tracking should be the desired goal of the system. If the ankle tracks position perfectly, it loses springlike behavior, which could be uncomfortable for the user, especially if they are still learning how to accurately control the wrist exoskeleton. In the torque control condition, we did not see this oscillatory behavior.

This technology has the potential to improve functional gait metrics by providing users with non-invasive sensory feedback and direct control of their prostheses, but the approach has practical limitations. One issue is that the user must attend to their wrist and cannot use their hand normally while walking with the device. If the benefits of direct control and sensory feedback were great enough, they might outweigh this cost and make a device using this approach viable on balance. It might also be that sensory feedback alone could be sufficient to improve gait, which would result in lower practical overhead; sensory information could be delivered by a smaller device that allows the user to move their wrist normally while walking.

Long term, we aim to use this system to test what users want from their prosthesis. Parameters for active prosthesis control have typically been hand-tuned to a generic control mode intended to work for an average user. However, customization using methods such as human-in-the-loop optimization (HILO) can substantially improve the efficacy of assistive devices [20]. We expect the same to be true for prostheses, but have not yet been successful, perhaps because the user has little sensory feedback to inform how they should best take advantage of each control law presented by the optimization system. We plan to test this system with HILO to determine whether the outcomes for functional gait metrics such as metabolic cost can be improved. In addition, because humans have been shown to continuously optimize metabolic cost [52], it is possible that the user could generate beneficial ankle trajectories with their wrist that are vastly different than those applied here, which were based on movements of the biological ankle.

There are many other scientific questions this novel teleoperation system could be used to address. For example, are people best able to operate the wrist exoskeleton with their dominant or non-dominant hand? Or, is it easier to learn using the wrist ipsilateral or contralateral to the amputation? Future work will address these questions. Systems like this could also be expanded in the future to incorporate an additional degree of freedom for medio-lateral stability, or untethered versions built to test for potential benefits during overground walking.

## VII. Conclusion

Our system closes the loop on both the control and sensory feedback from a robotic ankle-foot prosthesis via a novel wrist exoskeleton and teleoperation scheme. Benchtop tests of all system components confirm sufficient accuracy and responsiveness. We also demonstrate the feasibility of the system by confirming that a subject with a transtibial amputation can volitionally control the ankle prosthesis in different ways while walking, and that the system can control ankle prosthesis position accurately under these conditions. Future work will further examine this system with additional participants and examine its effects on functional gait metrics such as metabolic cost, phantom limb pain, and balance.

## Acknowledgment

The authors would like to thank our participant for his time and Susan Stenman, CP, for assisting us in fitting the prosthesis to our participant. We also thank Scott Delp for helpful insights and high-level feedback, as well as Laura Blumenschein, Margaret Koehler, Cole Simpson, and other members of the Collaborative Haptics and Robotics in Medicine Lab and the Stanford Biomechatronics Lab for their help with debugging, statistics, and editing. This work was funded by a National Science Foundation Graduate Research Fellowship to CGW (DGE-1656518) and National Science Foundation Grants CBET-1511177 and CMMI-1734449.

